# New imaging tools reveal live cellular collagen secretion, fibril dynamics and network organisation

**DOI:** 10.1101/2024.08.15.608104

**Authors:** Olivia Kent, Ellie R. Casey, Max Brown, Steven Bell, Matt Ehrman, Mike Flagler, Arto Määttä, Adam M. Benham, Timothy J. Hawkins

**Author notes:** Correspondence: Tim Hawkins Department of Biosciences Durham University main figures 3 supplementary figures 17 supplementary movies.

## Abstract

Although light microscopy has been used to examine the early trafficking of collagen within the cell, much of our understanding of the detailed organisation of cell deposited collagen is from static electron microscopy studies. To understand the dynamics of live cell collagen deposition and fibril organisation, we generated a bright photostable mNGCol1α2 fusion protein and employed a range of microscopy techniques to follow its intracellular transport and elucidate extracellular fibril formation. Our findings reveal the dynamics of fibril growth and the dynamic nature of collagen network interactions at the cellular level. Notably we observed molecular events that build network organisation, including fibril bundling, bifurcation, directionality along existing fibrils, and looping/intertwining behaviours. Strikingly, mNGCol1α2 fluorescence intensity maxima can mark a fibril before another growing collagen fibril intersects at this location. Our study shows that N-terminal protease site is not an absolute requirement for collagen fibril incorporation and is, to our knowledge, the first time that cell-directed collagen fibrillogenesis has been visualised at high resolution in real time. The approach paves the way for assessing the dynamic organisation and assembly of collagen into the extracellular matrix in skin models and other tissues during health, ageing and disease.

## Introduction

Collagens are essential extracellular matrix (ECM) proteins which account for approximately 30% of the total protein mass of mammals (Ricard-Blum 2011^1^). These collagens are diverse, comprising 28 fibrillar and non-fibrillar proteins, expressed in connective tissues including the skin, bone, and cartilage (Nimni 1983, Ricard-Blum 2011^1^). Collagen proteostasis is sensitive to both oxidative and reductive stress (Carne et al 2019^2^)Disruption to healthy collagen structure or expression is a significant factor in fibrosis, osteoporosis and several genetic diseases (Myllyharju & Kivirikko 2004^3^, Ricard-Blum 2011^1^) but is also a natural consequence of skin aging. For example, the *COL1A1* type I collagen is significantly down-regulated with age across photo-exposed and non-photo-exposed skin sites (Kimball et. Al. 2017^4^). Type I collagen, which accounts for 90% of human collagen, is comprised of two α1chains (expressed by the *COL1A1* gene) and one α2 chain (expressed by the *COL1A2* gene) that associate as a triple helix heterotrimer to form a procollagen polypeptide of approximately 300 nm in length (Canty & Kadler, 2005^5^). This heterotrimer assembles in the ER under the guidance of multi-functional chaperones, such as PDI (P4HB), its partner in the prolyl-4-hydroxylase complex, P4HA, and the collagen-specific chaperone Hsp47 (Xiong ^6^ et al 2018, Wilson ^7^ 1998, Ishida and Nigata 2011 ^8^). Procollagen undergoes extensive processing before and after secretion from the cell, including N- and C-terminal proteolytic processing and fibrillogenesis (Tanaka 2022^9^, Kelly, 1985) Although collagen egress from the ER and ER-Golgi transport (Brown et al. 2014^10^) has been extensively studied, less is known about the post-Golgi transport of collagens to the plasma membrane (PM) and the rate determining steps of collagen secretion in different cell types. Following secretion, collagen is deposited into the ECM where, facilitated by fibronectin, it self-assembles into a fibrous network to provide connective tissues with mechanical strength (Birk et al. 1989, Kadler et al. 2000).

Although electron microscopy can inform us about the complex organisation of preassembled collagen networks surrounding cells and tissues, dynamic information about how collagen fibrils elongate and are organised by living cells to generate these networks is limited. By pairing a novel bright and stable collagen fusion protein with advanced high temporal and spatial microscopy techniques, we have charted the trafficking and assembly of collagen, capturing individual secretion events and observing the live-cell dynamics of collagen deposition, fibril growth and organisation from live cells. Overall, the study reveals how dynamic phenomena establish network organisation at the cellular level.

## Results

### Construction of an mNG-type I collagen fusion protein marker, mNGCol1α2

To follow the trafficking, exocytosis and fate of type I collagen at high spatial and temporal resolution we created a fusion protein combining the bright and stable fluorescent protein mNeonGreen (mNG) with the Col1α2 chain. mNG, a green/yellow fluorescent protein isolated from the marine invertebrate *Branchiostoma. lanceolatum,* is ∼4 fold brighter and more photostable than GFP, allowing it to be detected by lower laser intensities, over longer periods and a higher frame rates (Shaner et al., 2013^11^; Tanida-Miyake et al., 2018^12^, Hostettler et al., 2017^13^). A similar fusion protein strategy, employing an N-terminal GFP instead of mNG, has been used to observe collagen *in vitro* (Lu et al 2018^14^), and at the tissue level in a zebrafish model (Morris et al., 2018^15^). Tagging Col1α2 is preferable to Col1α1 because two tagged Col1α1 monomers within the heterotrimeric complex could increase the possibility of steric effects and produce variation in labelling intensity. The fusion protein in this study retains the N terminal signal peptide, replacing the N-propeptide (pp)with mNG, which is flanked by GS linkers to confer flexibility **[Figure 1A]**. Importantly, removal of the N terminal ADAM proteinase site prevents proteolysis of the mNG tag on collagen following secretion of the fusion protein into the ECM enabling the observation of fibrillogenesis.

**Figure 1:**
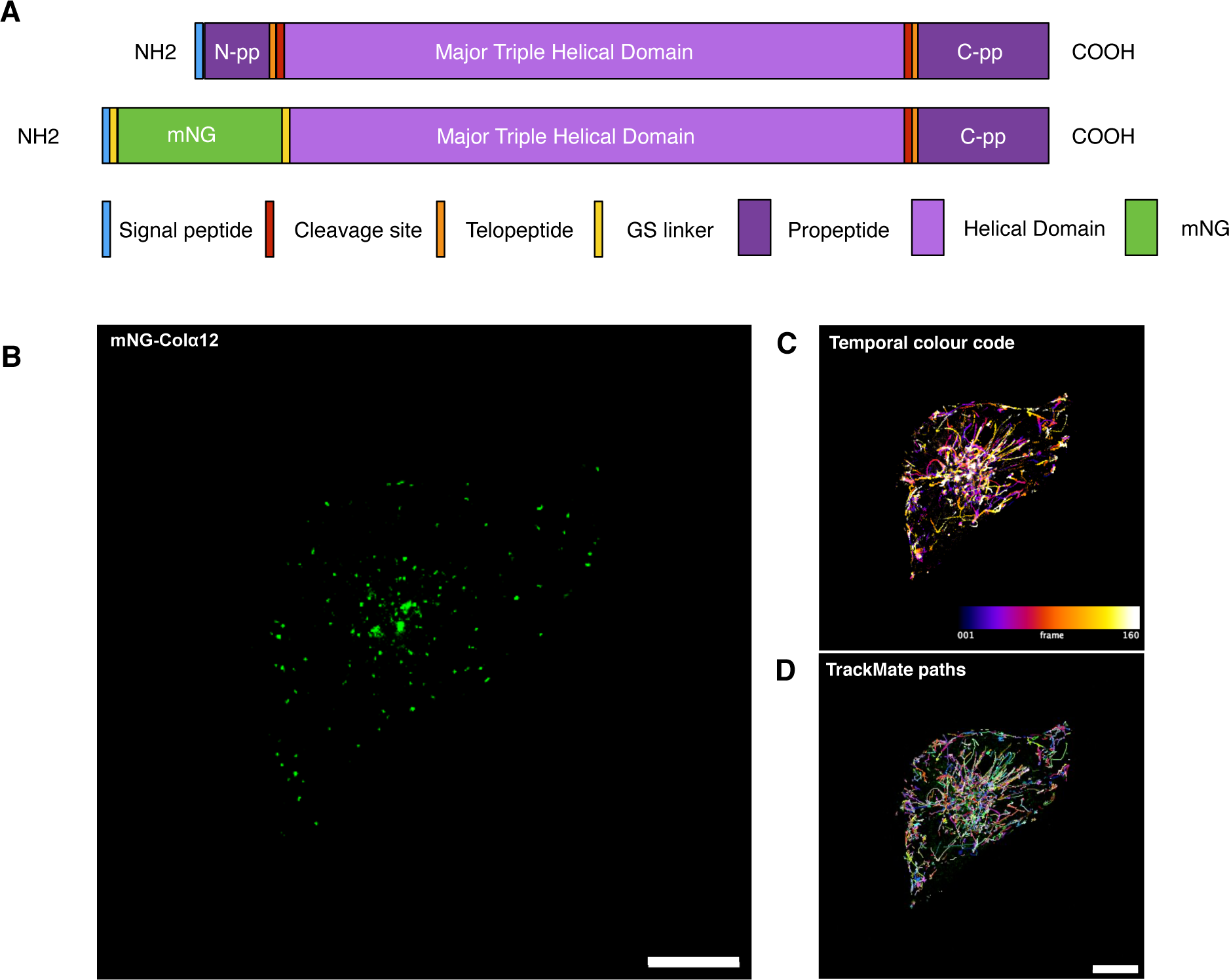
Live cell imaging of mNGCol1α2 expressed in HT 1080 cells. **(A)** Diagram of mNGCol1α2 fusion protein. **(B)** HT1080 cells expressing mNGCol1α2. mNGCol1α2 marks small rapidly moving particles which predominantly move towards the periphery of the cell. **(C)** Time coding projection illustrates passage of particles towards the periphery. **(D)** Particles were isolated and tracked using TrackMate and Fiji to calculate size, number and velocity. Scale = 10μm, 206.44 ms per frame or 4.844 fps

The mNG-Col1*α*2 fusion was tested first in HT1080 fibrosarcoma cells, which correctly assemble and quality-control collagens but lack endogenous expression of Col1, enabling mNGCol1α2 to be evaluated in the absence of competitor Col1 chains.

### mNG-Col1α2 traffics from the ER, interacts with expected folding partners and becomes hydroxylated

To confirm that mNGCol1α2 was correctly processed and post-translationally modified, mNG-Col1a2 was immunoprecipitated (IP) from transfected HT1080 cells, trypsinised and subjected to data-dependent acquisition mass spectrometry (DDA MS). mNG-Col1α2 interacted with key processing proteins including P4HA and P4HB (PDI) **[Supp Figure 1A]**, (Rappu, Salo, Myllyharju, et al. 2019^16^). To confirm the integrity of the fusion protein, the peptide sequences of the trypsin-digested immunoprecipitates were assessed by protein pilot. Peptide coverage spanned the entire fusion protein, and mNG-Col1α2 was hydroxylated at multiple sites **[Supp Figure 1B]**, demonstrating that the fusion protein could receive specific post-translational modifications in HT1080 cells

To confirm that the mNGCol1α2 trafficked, the exit of proteins from the ER was prevented by treatment with Brefeldin A (BFA). Prior to treatment, mNGCol1α2 was distributed between the ER, a post-ER compartment likely to be the ERGIC, and small vesicles **[Supp Figure 1C]**. Following treatment, high levels of colocalization of mNGCol1α2 with the ER resident protein PDI (P4HB) in the ER were observed, indicating retention. **[Supp Figure 1D]**. A statistically significant increase in colocalization was quantified using the Coloc2 plugin for Fiji. BFA treated cells show a Pearson’s R value of 0.96 compared to 0.26 for control cells, indicating significant colocalization between mNGCol1α2 and PDI **[Supp Figure 1D]**. Following BFA treatment, peripheral vesicles were lost, consistent with the expectation that newly synthesised collagen is trafficked from the ER to these carrier vesicles.

### mNGCol1α2 is packaged into highly dynamic collagen carriers

In transfected HT1080 cells observed at “steady-state”, the mNGCol1α2 fusion protein was localised to numerous punctae within the cytoplasm consistent with vesicles **[Figure 1B]**. These carriers exhibited two distinct morphologies, either spherical (60-80%) or tubular (20-40%). The diameter of spherical carriers ranged between 358 nm and 748 nm with a mean value of 510 nm. The peripheral location of these structures suggests that they represent Golgi to PM carriers (GPCs). The collagen carriers were highly dynamic with movement principally towards the plasma membrane, although objects moving away from the PM were also observed **[Supp movie 1]**. Temporal colour coding **[Figure 1C]** was used to demonstrate the trajectory of vesicles with the white latter frames showing vesicles accumulating at the periphery of the cell, often within cellular projections. Motile vesicles were tracked in time-lapse movies using i ImageJ and TrackMate. A representative dataset with particle tracking is shown in **[Figure 1D and supplementary movie 2].** Within just 60 seconds, the observed particles have transversed the cell, with a mean velocity of 0.83 μms^-1^ (SD) 0.14 μms^-1^. The enhanced mNG signal intensity allowed short exposures and high frame rates (10-206ms per frame) enabling accurate capturing of rapid movement without blurring or gaps in the vesicles path **[Figure 1C, Supplementary movie 1]**.

### Delivery of mNGCol1α2 collagen carriers to PM requires microtubules

To further investigate the dynamics of collagen carriers, we studied the microtubule dependency of vesicle movements. Early studies reported that disruption of microtubules by colchicine impairs collagen secretion from fibroblasts (Diegelmann and Peterkofsky 1975^17^). Co-labelling of microtubules in HT1080 cells expressing mNGCol1α2 revealed that the majority of collagen carriers associate with microtubules **[Figure 2A & B, Supp movie 3].** High speed super-resolution Airyscan confocal microscopy allowed these small but well-defined carriers to be seen clearly, tracking along the microtubule **[Supplementary movie 3**].

**Figure 2:**
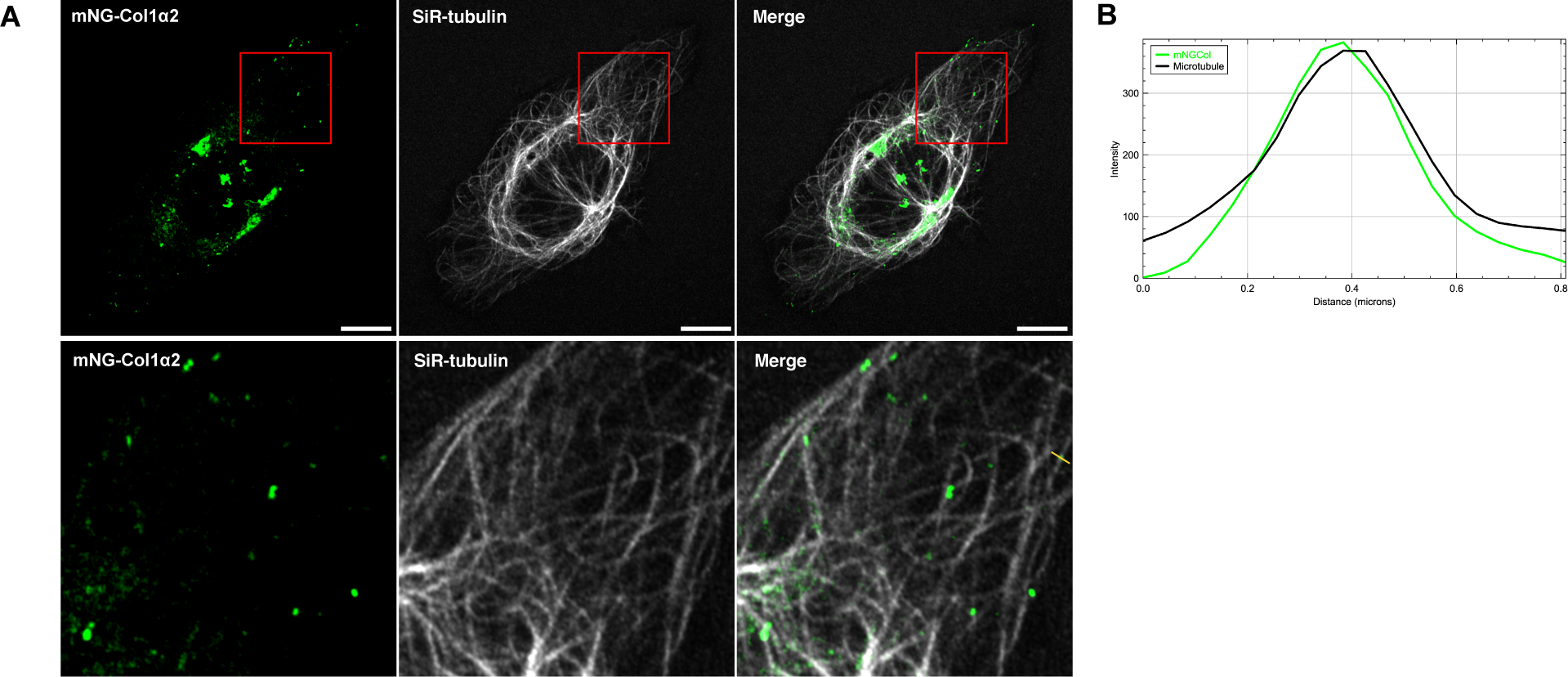
mNG-Col1α2 containing vesicles associate with and move along the microtubule network. **(A)** Live cell imaging of mNGCol1α2 with SIR-tubulin in HT1080 cells. mNGCol1α2 particles align along microtubules and move along these paths. **(B)** Quantitative line profile of co-localization along the yellow line in A. Scale = 10 nm.

### High speed TIRF microscopy reveals the fate of collagen containing vesicles

Having observed trafficking of mNGCol1α2 through the secretory pathway towards the PM, we used mNGCol1α2 to study the less understood process of collagen exocytosis. Total Internal Reflection Fluorescence Microscopy (TIRF) produces a thin <200nm excitation field at the glass water interface allowing real time visualisation of objects at the PM. The technique has been used to provide insights into the movement of organelles and single molecules at the PM including the moment of exocytosis of synaptic vesicles containing neurotransmitters into the synaptic cleft (Midorikawa, 2018^18^). We used ring TIRF microscopy to follow the vesicles at high temporal resolution (10-100 ms). Multiple bursts occurred at the PM within the 200 nm TIRF field which represent the moment of exocytosis. Bursts mostly occurred at the Baso-lateral surface of the cell predominantly towards the lateral periphery **[Figure 3A]**. Motile carriers approaching the PM stopped, becoming static for a short period, before ‘bursting’ and releasing the mNGCol1α2 cargo **[Figure 3B]**. The high frame rate resolves the release and subsequent dispersal of the fluorescent signal **[Figure 3B**, **Figure 3C and Supp movie 4]**. Observed pausing events of a mean duration of 30 seconds may represent the capture of the carrier by the actin/myosin cytoskeleton at the PM [**Figure 3D].** Intriguingly in the example shown in **[Supplementary movie 4]** with 10ms temporal resolution, the carrier appears to transition past an exocytosis location and return, before becoming stationary and releasing its contents.

**Figure 3:**
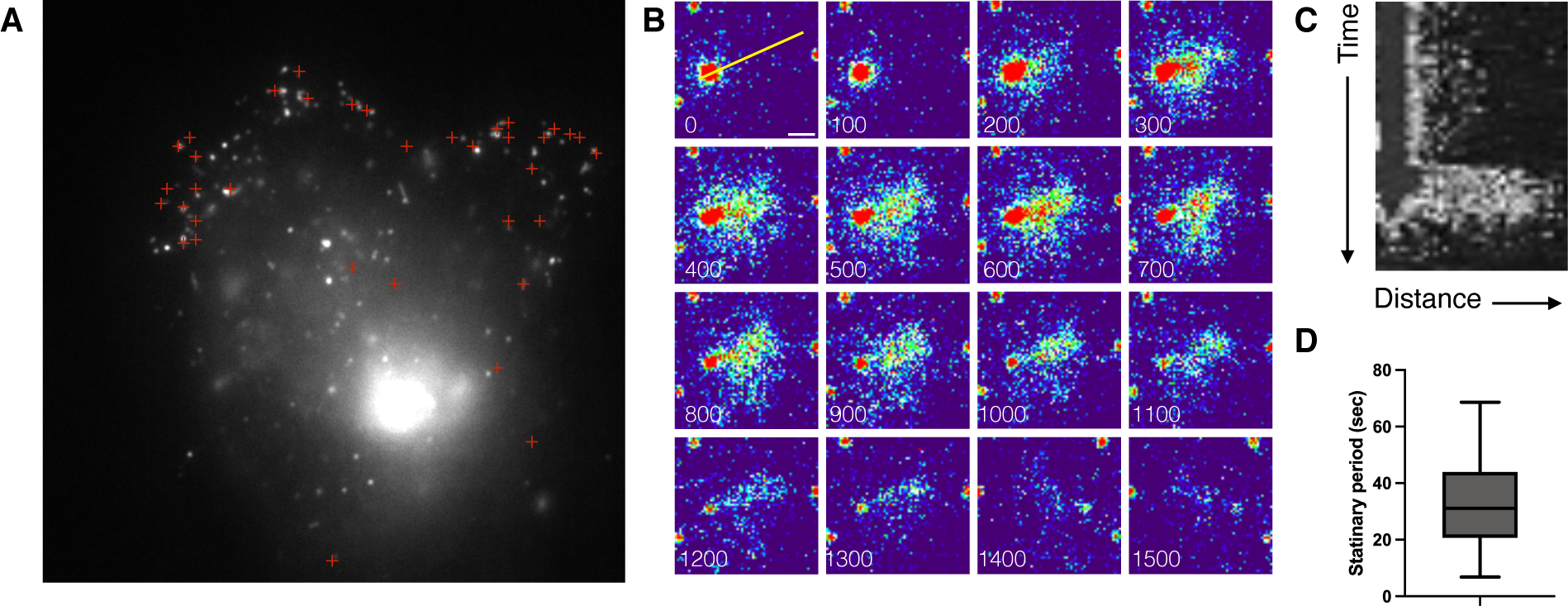
mNGCol1α2 secretion events. **(A)** Full field of view TIRF image of an mNGCol1α2 expressing HT1080 cell. Red + mark locations of collagen ‘burst’ events. These events usually clustered at the periphery of the cell on the lower surface. **(B)** TIRF movie frames of a burst event. 100ms between frames. Scale = 1μm **(C)** Kymograph of burst event derived from yellow line in B. **(D)** Box and whisker plot of stationary phase of vesicles at the PM before they fuse with PM to give a burst event.

### Super-resolution imaging of extracellular deposited mNGcollagen fibrils from live cells

To investigate the process of collagen fibrillogenesis and deposition of fibrils within the ECM we turned to Saos-2 osteosarcoma cells expressing mNGCol1α2. Unlike HT1080, Saos-2 cells express both type 1 Col1α1 and α2 chains, and highly express the lysyl hydroxylase LH1 to support the extra-cellular cross-linking of collagen (Fernandes et al. 2007^19^). Transfected Saos-2 cells produce labelled collagen carrier vesicles with a velocity of 0.79 μms^-1^ (SD 0.14 μms^-1^) **[Figure 4A, Supp Movie 5]**, comparable to those seen in HT1080 cell. Furthermore, these vesicles also move along microtubules, demonstrating that type 1 collagen produced by different cell types utilises the same post-ER transport machinery **[Figure 4B and Supp movie 6 and 7].** TrackMate analysis was used to chart these vesicle movements along microtubule tracks. As for HT1080 cells, BFA treatment of Saos-2 cells caused the loss of collagen carriers and accumulation of the collagen signal within the ER **[Supp Figure 2].** To examine the biosynthesis of collagen fibrils outside the cell, Saos-2 cells were cultured in the presence of ascorbic acid for 48 hrs on fibronectin-coated coverglass-bottomed dishes and imaged with super-resolution microscopy (Airyscan and 3D-SIM). To date, dynamic super-resolution imaging of deposited collagen has not been documented in the published literature. This experiment revealed the first dynamic observation of deposited collagen from live cells at 100 nm resolution and clarity. Fibrils containing mNGCol1α2 were deposited beneath the cell, observed to be linked to the cell, or as networks close to a cell. When imaged with Airyscan with JDCV processing, the collagen fibrils had a mean Full Width Half Maximum (FWHM) of 106 nm (range 91-131 nm) consistent with observations of collagen fibril diameters of 35-110 nm in electron microscopy studies (Parry et al 1978^20^, Parry and Craig 1984^21^) This diameter can vary depending on tissue up to approximately ∼ 300nm **[Figure 5A & 5B].** These mNG-Col1α2 containing fibrils were readily detected by an antibody that recognizes native triple helical type I collagen **[Figure5A].** Both straight and curved fibrils were within the size range expected. Importantly, these results show the integration of mNGCol1α2 into the lattice of unlabeled native collagen to generate a singular fibril or bundle **[Figure5A merge insert**]. Of note, 3D-SIM super-resolution microscopy resolved small loops within individual fibrils in 3D **[Figure 5C]**. 3D-SIM bleaching experiments showed that the mNG signal did not diminish equally throughout the length of the fibril **[Supp Movie 8].** The distribution of the mNG-Col1α2 bleaching may further reflect the number and fine spatial incorporation of the fusion protein with the collagen fibril.

**Figure 4:**
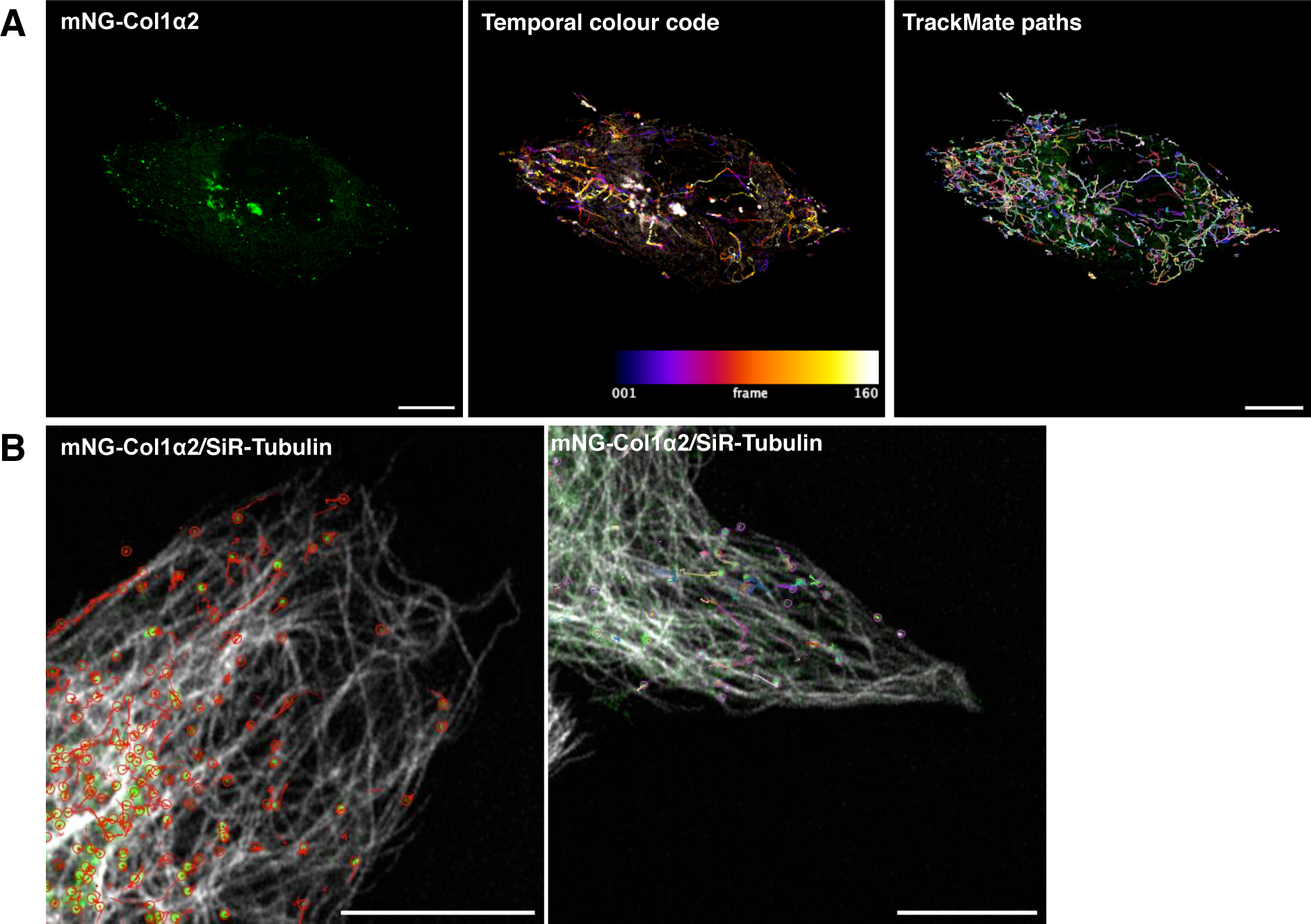
mNGCol1alpha2 labels fast-moving carriers in Saos-2 cells with movement along microtubules. **(A)** mNGCol1α2 labels collagen carriers or vesicles which rapidly move towards the periphery. These paths are shown by temporal colour coding. TrackMate tracks are used to quantify vesicle size and velocity Scale 10um. **(B)** Saos-2 cells expressing mNGCol1α2 co-imaged with microtubules labelled by SiR-tubulin. Overlay shows mNGCol1α2 association with microtubule. Superimposed on these images are the paths of TrackMate detected paths of mNG-Col1α2 carriers demonstrating their movement along microtubules.

**Figure 5:**
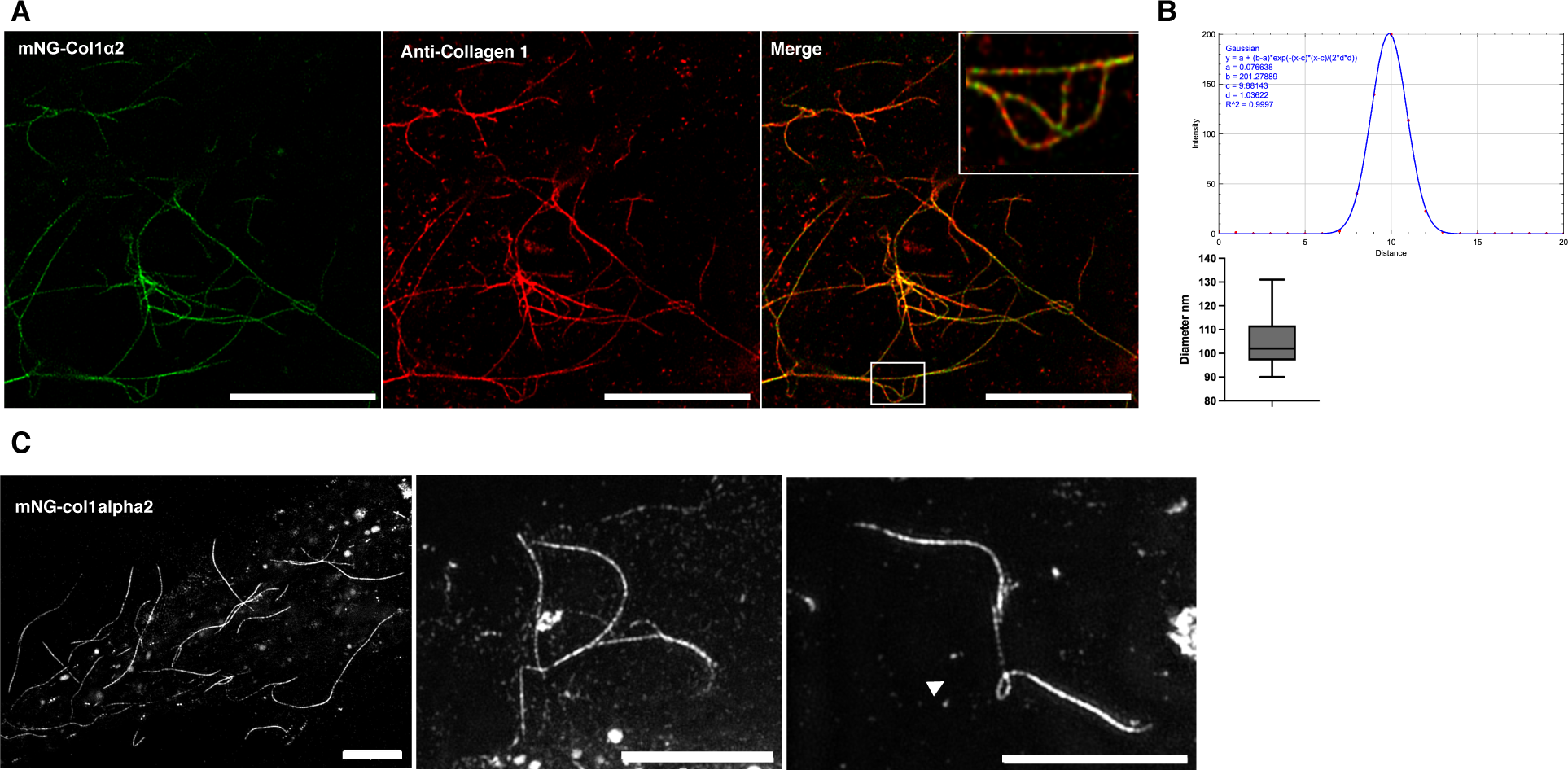
Superresolution microscopy of collagen fibrils containing mNG-Col1α2 deposited by Saos-2 cells. **(A)** Airyscan jDCV Super-resolution image of deposited mNGCol1α2 collagen fibrils (Green). Fibrils are co-labelled with anti-Collagen 1 antibody (red). Merge - yellow shows areas of colocalization along the fibril. Zoomed area shows the relationship between accessible native Col1 epitopes and the incorporated mNG-Col1α2. Scale = 10 μm **(B)** FWHM analysis of deposited collagen in A. Box and whisker plot shows range and mean of collagen diameter values. **(C)** 3DSIM super-resolution imaging of deposited mNG-Col1α2 fibrils. These three-dimensional stacks reveal loops within fibrils. Scale = 5 μm

To establish the utility of this platform to compare collagen trafficking and fibril growth across collagen producing cell types we also confirmed that comparable vesicular carriers and fibril deposition was observed in BJ fibroblast cells expressing mNGCol1α2 **[Supplementary** Figure 3]

## Extracellular collagen fibril dynamics

The deposited collagen network was dynamic even over short timescalesof minutes where it was seen to flex, potentially due to new fibril incorporation and attachment or the motility of the overlying cell. Longer term live cell imaging of mNGCol1α2 over 18-20 hrs enabled the observation of extracellular collagen fibril growth and dynamics in unprecedented detail and revealed the existence of a range of fibril dynamic phenomena which have to date not been reported. The directional growth of an individual collagen fibril was tracked and found to have an average rate of 0.11 μmin^-1^, with these often radiating from a preexisting area of bundled fibrils **[Figure 6A and Supp movie 9].** Fibrils followed straight and curved trajectories with periods of both slightly slower and faster growth rates, including pauses, but maintaining a positive trajectory **[Figure 6B]**. In our experiments negative trajectories, which could potentially indicate filament breakdown or turnover by MMPs, were not observed. In addition, we observed at least six distinct dynamic process or organisational phenomena:

1. **Growth of fibrils along pre-existing fibrils with bundling and bundle bifurcation** –additional fibrils grow along existing fibrils (black arrow follows the tip of this second fibril) **[Figure 7A, Supp Movie 10]**. The fibrils increased in intensity and widened as new fibrils extend along the exiting network, likely indicating packing together and bundling **[Figure 7B & C]**. Monitoring of fluorescence intensity within a defined ROI (red box) shows that the intensity of this zone increased in three steps representing each fibril passing through **[Figure 7D]**. We observed these bundles to bifurcate as the linked fibrils separated to follow different trajectories, or started to zipper and snap together. **[Figure 7E, Supp movie 11].**
2. **Fibril crossover –** collagen fibrils grew across existing fibrils, where the increase in collagen signal at nodes suggests collagen reinforcement of the site **[Figure 8A Arrows, & Supp movie 12]**. Strikingly, when movie frames are examined closely the maxima can form before the crossover occurs at that location (red circles), potentially predetermining the site of interaction **[Figure 8B]**. In some cases, a puncta of collagen forms on an existing fibril and grows in intensity; the approaching fibril crosses at this point and further intensification at this location follows the crossover event. **[Supp Movie 9].**
3. **Meeting and joining** – fibrils growing in opposite directions can meet and join. In these circumstances, the two fibrils appear to move closely past each other and join potentially through bundling to form a new thinker and singular bundle. **[Figure 9A and Supp Movie 13]**.
4. **Tensioning** - curved and flexible fibrils can tension into angular arrays, likely driven by mechanical forces exerted by the cell above **[Figure 9B and Supp Movie14].**
5. **Looping**-fibrils often looped back on themselves during growth **[Figure 9 C and Supp Movie 15],** or pre-existing fibrils were looped or pushed up by mechanical forces exerted by the cell. This looped fibril architecture was clearly resolved with 3D-SIM **[Figure 9D, Supp Movie 16].** Once growing fibril loops are formed, they can be fortified further with collagen as new fibrils thicken the loops. **[Figure 9C and Supp Movie 15]**.
6. **Intertwining** – as well as bundling individual fibrils or bundles of fibrils can wind around each other with crossovers potentially forming the basis for a larger and thicker accumulation of collagen or network. **[Figure 9E and Supp Movie 17]**. The multiple ways that collagen fibrils can grow and become organised is summarized in **Figure 10**.

**Figure 6:**
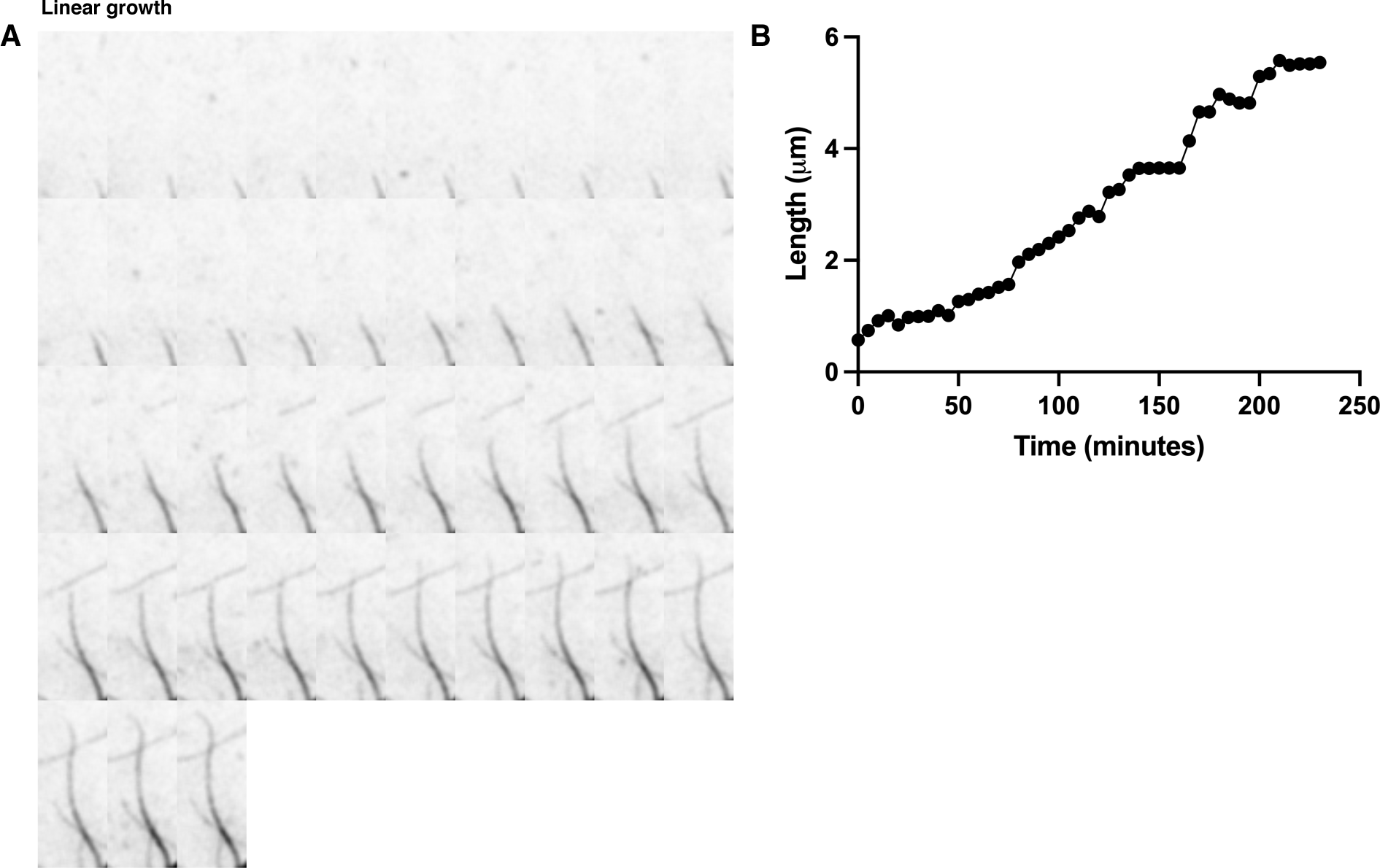
mNG-Col1α2 fibrils exhibit sustained directional linear growth. Frames from a Saos2 mNG-Col1α2 fibril deposition movie (subsection) showing linear growth of a fibril being tracked. **(B)** Graph shows the length of an isolated collagen fibril over time showing the rate of elongation with some pauses and or moments of stalling.. Frame rate is 1 frame every 5 minutes or 12 frames per hour.

**Figure 7:**
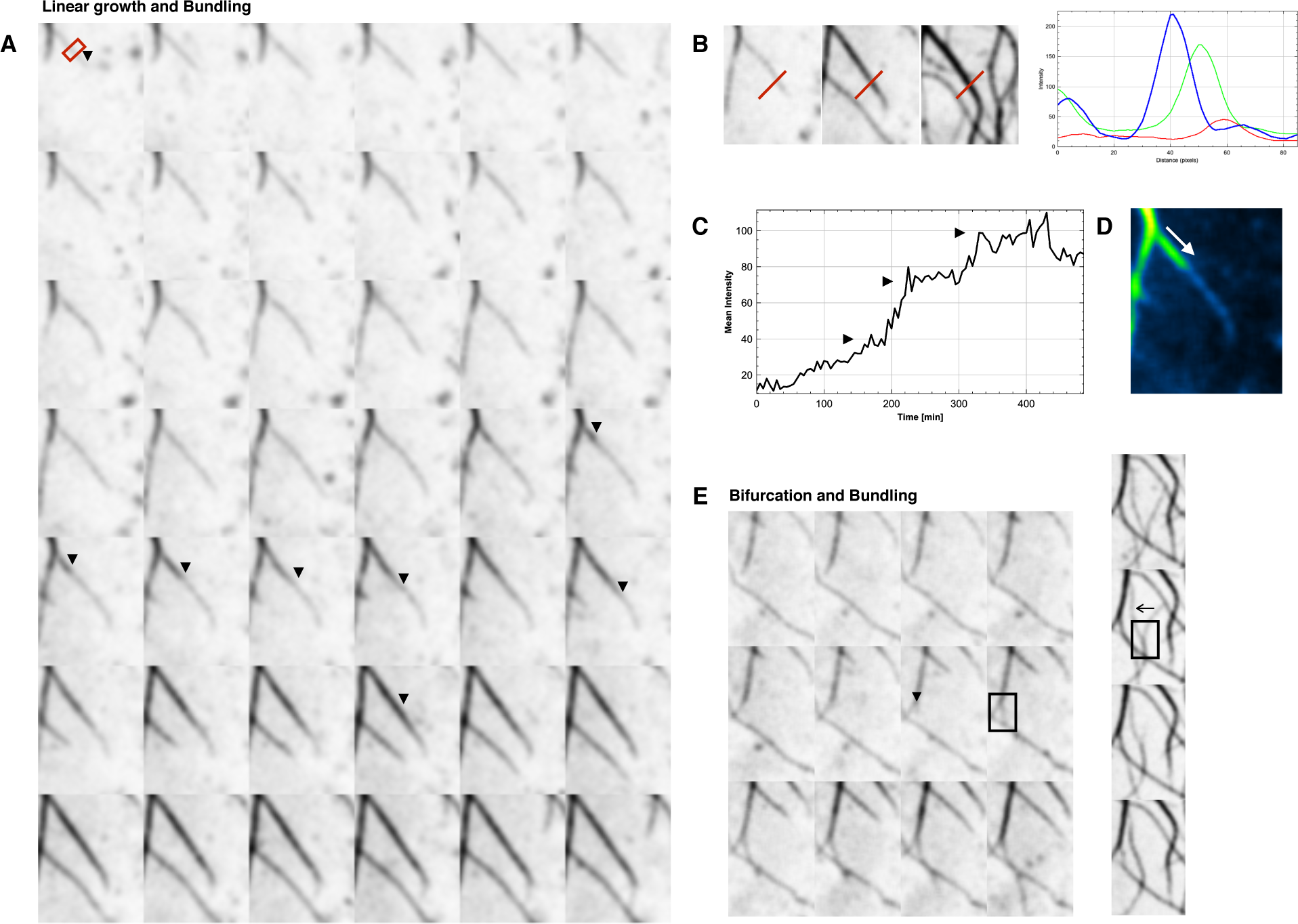
Collagen fibrils can grow along or coaligned with existing fibrils. **(A)** Frames from **an** mNGCol1α2 fibril deposition movie showing growth of collagen fibrils along existing fibrils with potential bundling and increase in signal intensity. **(B)** Three frames taken from movie in A demonstrating three events of “on fibril” or coaligned fibril growth and bundling. Line plots of red line show the intensity and width of the fibril increasing. **(C)** Time intensity plot of the red box zone in A shows the three events when a fibril travels along the underlying existing fibril. Events marked with black arrow. **(D)**The direction increase in fibril intensity as a second fibril growth along another is illustrated with the use of a LUT. **(E)** Frames from an mNGCol1α2 fibril deposition movie showing a bifurcation event. Site of bifurcation highlighted by black arrow. The bifurcation event occurs within the black box. Frame rate is 1 frame every 5 minutes or 12 frames per hour.

**Figure 8:**
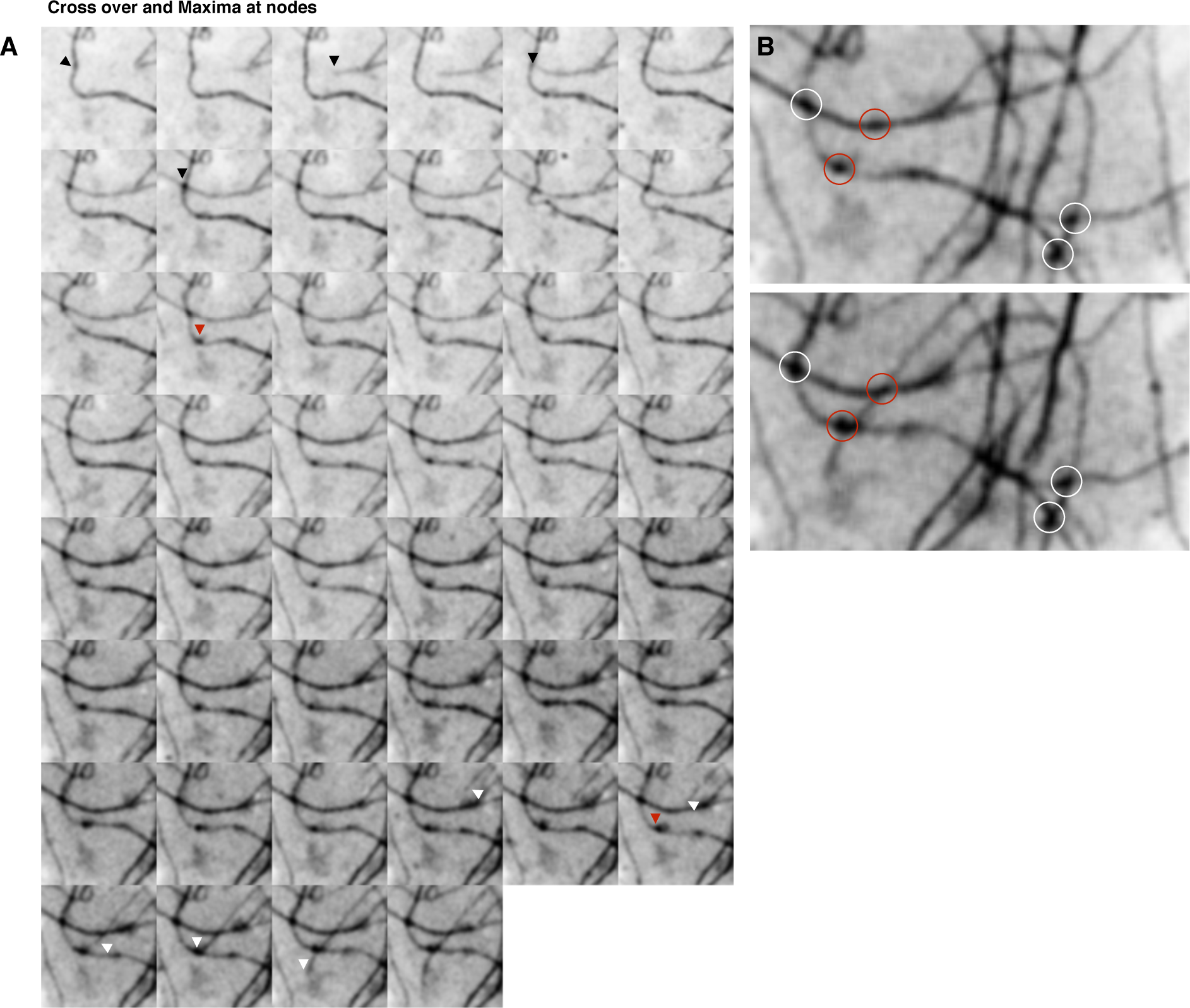
Locations at which collagen fibrils cross over have distinct mNG-Col1α 2 fluorescence intensity maxima which can exist before a crossover event. **(A)** Frames from an mNGCol1α2 fibril deposition movie. Points of fibril-fibril crossover locally increase in intensity to give dark foci representing a local accumulation mNGCol1α2. 5 minutes per frame. **(B)** Dark foci of mNGCol1α2 accumulation can pre-exist along a fibril. A new growing fibril entering the frame can be seen to transition specifically through these points (Red circles) suggesting a predetermination of the location of these crossover nodes. Frame rate is 1 frame every 5 minutes or 12 frames per hour.

**Figure 9:**
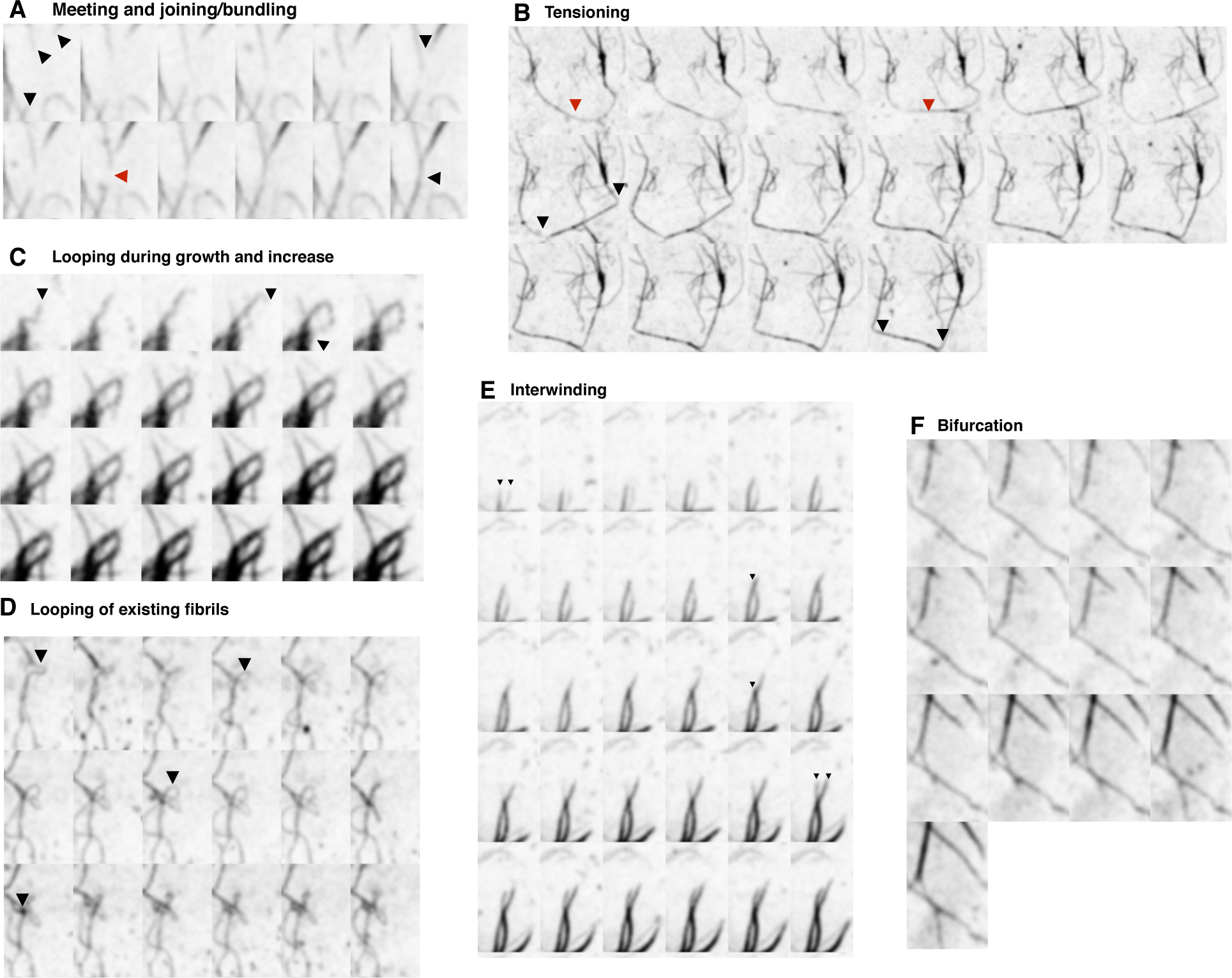
A wide range of further fibril interactions and organizational phenomena are observed. **(A)** Frames from a mNG Col1α2 fibril deposition movie. Fibrils highlighted by black arrows grow towards each other before joining at the red arrow. **(B)** A fibril marked by a red arrow straightens under tension. Two corner attachment points are marked by black arrows. **(C)** Growing fibrils produce a loop structure that is then further strengthened by addition of more mNGCol1alpha2 to give a thick loop **(D)** Existing fibrils can be pulled into loops by flow or mechanical force from the cell above. Both of these loops could represent further sites of attachment for other motile cells **(E)** Frames from a mNG Col1α2 fibril deposition movie. Two separate mNGCol1α2 labeled fibrils interwine around each other followed by an increase in fibril intensity and width. Frame rate is 1 frame every 5 minutes or 12 frames per hour.

**Figure 10:**
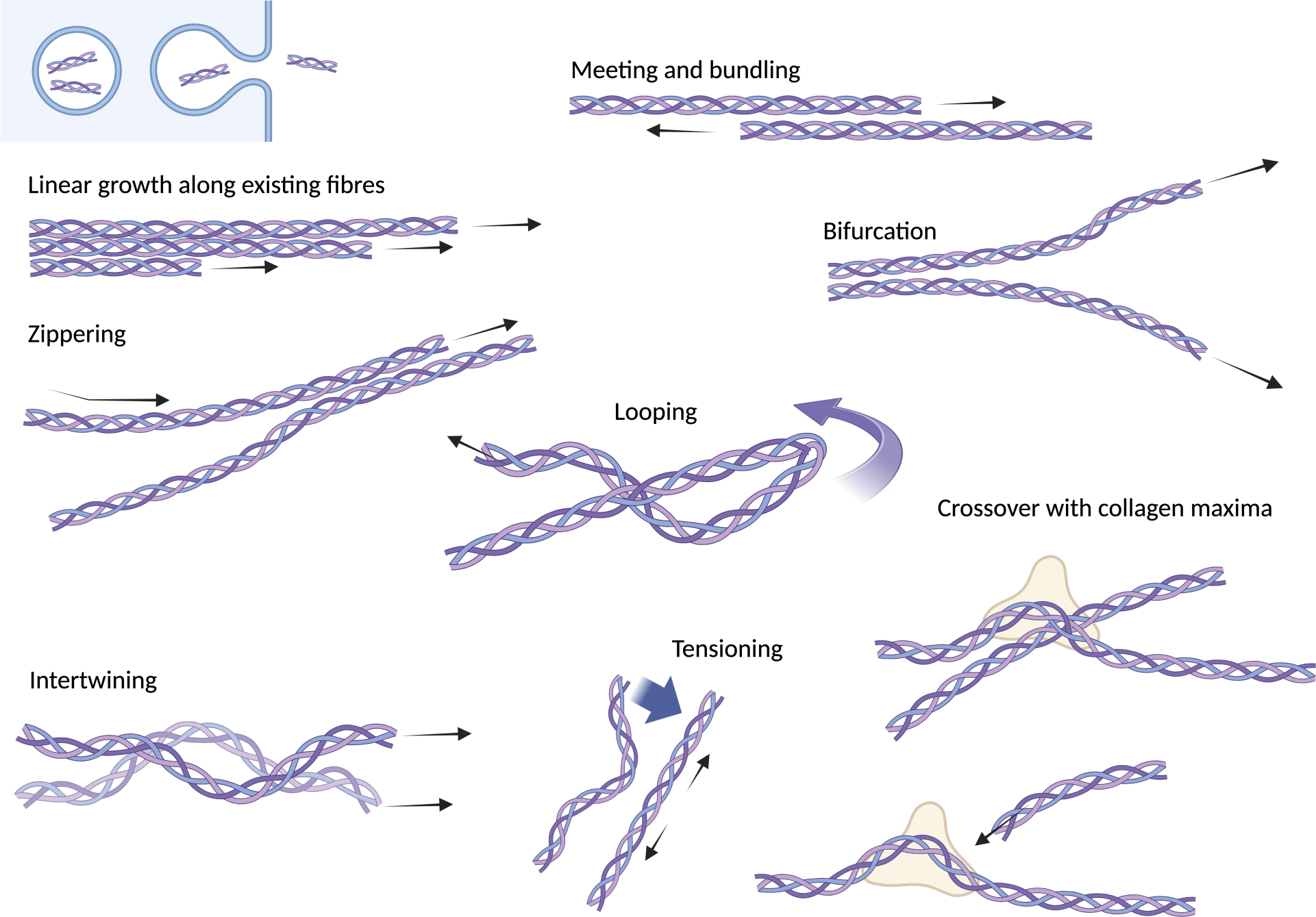
Summary diagram of the observed multiple ways that collagen fibrils can grow and become organised.

## Discussion

In this study, we have captured the sought-after moment of secretion of a collagen molecule with high temporal resolution and have developed a system for imaging live cell fibrilogenesis and organisation for the first time.

The mechanisms by which cells direct complex packages of collagen to the ECM remain largely unknown. We show that rapidly moving collagen containing packages pass along microtubules towards the cell periphery, consistent with the kinesin driven collagen trafficking in myofibroblasts (Kamata 2017^22^). The collagen carriers are captured at the cell membrane, remaining stationary for ∼ 30 seconds until they fuse with the PM and release their cargo. Further co-labelling studies will be required to identify the molecular mechanism involved, but it is likely to require actin capture or a SNARE based exocytosis processes.

Alternatively, additional motor proteins, such as myosins, could be involved to facilitate short distance movement, membrane fusion and secretion. Collagen containing packages may be captured by myosin association with actin at the PM as seen in melanosomes exocytosis (Gross 2002^23^). Notably, within our system, secretion sites and the location of fibril nucleation/growth appear to arise independently.

We show that, following secretion from Saos-2 cells, the mNGCol1α2 fusion protein can produce extra-cellular fibrils and successfully incorporate native type 1 collagen into these fibril arrays, with superresolution microscopy capable of visualising these sites along a fibril. Our live cell imaging of fluorescent collagen in Saos-2 cells revealed the dynamic nature of collagen assembly, with the fibrils exhibiting a range of organisational interactions. Several of these observed phenomena have been predicted from electron microscopy of collagen matrices in tissue, yet this is the first observation of these higher order networks occurring and developing in a living cell at the subcellular level. Processing and fibril formation is considered to occur outside the cell through the secretion of procollagen with fibronectin also facilitating procollagen cleavage at the cell surface (Saunders & Schwarzbauser 2019^24^). We observed the formation of fibrils on the underside of the cell in agreement with studies that suggest that fibril assembly is likely a cell-controlled process occurring at the plasma membrane where the cell surface assists in collagen assembly (Kadler et al 2008^25^ Curr. Opin. Cell Biol., Velling et al J. Biol. Chem. 2002^26^.) Whether this surface collagen assembly is facilitated by fibronectin interactions, or integrins directly is debated (Kadler et al 2008., Velling et al J. Biol. Chem. 2002^26^, Musiime et al 2021^27^).

The mNGCol1α2 containing fibrils in our experiments were readily detected by antibodies that recognise native triple helical collagen and grow at a rate of several μm per hour and often, but not exclusively, radiate from locations of existing dense collagen accumulations. Some static electron tomography studies have observed fibrils extending from membrane invaginations and projections called fibripositors (Kalson et al 2013^28^). The authors show that following secretion, fibrils are formed at the cell surface by nucleation of collagen, just as we observe **[Figure 5A & C],** The newly formed fibrils may subsequently be pulled into a membrane recess by myosin (NMII) at their membrane anchors. At this point, fibrils would continue to grow by molecular accretion of secreted collagen, whilst tension is continually exerted by NMII. The same EM study identified that fragmented or unanchored fibrils can be totally internalised in a process requiring both NMII and dynamin to form fibricarriers (Kalson et al 2013^28^). Fibripositors have only been observed in embryonic tendon cells where linear organisation of collagen is required, with ssTEM or SBF-SEM and were not observed in our live system. It will be of interest to translate our findings to tendon cells to determine the dynamics of fibripositor formation in the future.

We are the first to observe growing fibrils interacting with each other beneath a live cell, joining tip to tip or meeting and then growing in parallel zippering-type activity **[Figure7].** Furthermore, bundles can bifurcate or yield Y shape organization. In agreement with the appearance of fibrils in our study, this behavior has been observed in EM 3D reconstruction studies where fibrils have been seen to interact tip to tip, with the end of fibrils fusing together or through tip to shaft interactions (Kadler et al. 2000^29^, Starborg et al 2009^30^, 2013^31^). *In vitro* studies have also shown that ends of fibrils can fuse together (Kadler 2000) whilst examination of individual collagen fibrils from the skin of 12-week-old mice has identified branched networks of collagen fibrils (Kadler 2000). Together, EM data of native collagen in these papers verifies that the collagen fibril behaviors reported here are physiological and are not caused by steric hindrance of the fusion protein.

Collagen bundling and higher organisation is controlled by fibril-associated collagens FACITs such as Col V and Small Leucine Rich Proteoglycans (SLRPs) such as decorin expressed in Saos-2 cells (McQuillian et al 1995^32^). These SLRPs decorate fibrils and regulate and limit the diameter of fibril bundles in tissues via their glycosaminoglycan chains which wrap around the fibril (Ricard-Blum 2011^1^) They are also insulators that prevent fibril–fibril surface interactions which can lead to fusion (Graham et al 2000^33^). Knockouts of decorin or fibromodulin show increased branching of collagen networks (Kadler et al 2000^29^). We observed fibrils growing towards each other, sliding past and bundling, presumably creating an antiparallel fibril as well as parallel zippering **[Figure7E, 9A]**. EM analysis of tissues has shown that they can contain both polar and unipolar fibril bundles (Kadler 2000^29^). Importantly, we have observed dynamic joining together of fibrils that evolve as the network develops, enabling the critical long-range transduction of force.

The increase in mNG-collagen fluorescence at nodes or junction points to create distinct dark foci is interesting and may identify a previously unobserved structural reinforcing of crossover points of fibrils. Within this zone there is accumulation of collagen, potentially as short fibrils which interact with the existing fibril either directly or via other fibril-associated proteins. The composition of these nodes and their interactions with cells merit further study.

Cells can remodel existing collagen fibril networks around them to transmit forces and communicate with other cells by exerting stress on nearby fibrils driven by the actin/myosin cytoskeleton (Canty et al 2006^34^). In our timelapse movies, cells can exert force on an existing fibril, tensioning it. In **[Figure 9B]** the fibril appears to have two connection points with the cell which hold onto the fibril straightening it and creating right angle kinks. Such fibril to cell surface connections are likely to be those established by integrins which can bind directly to collagen such as α1β1 in this cell line (Vihinen et al 1996^35^) or indirectly via collagen-integrin bridging molecules such as fibronectin. (Zeltz et al 2014^36^, Bourgot et al 2020^37^, Zeltz et al 2016^38^, Zeltz et al 2020^39^). Local exertion of force onto fibronectin can promote fibronectin fibrillogenesis, consequently facilitating local collagen assembly at these sites (Musiime et al. 2021^27^). Which of these mechanisms are at play will depend on the integrin repertoire of different cell types.

In conclusion, coupling an mNGCol1α2 fusion protein with advanced imaging technologies has enabled us to reveal the molecular detail of collagen secretion and fibril production by living cells, providing novel and fundamental insights into the dynamic mechanisms for ECM assembly and organisation of the local collagen network. These observations shed light on how interactions of the newly secreted collagen link to both the cell and other collagen fibers, enable construction and regulation of network complexity at the cellular level. Translating this approach to different tissue types and collagen types will further our understanding of the importance of ECM composition, organisation and stiffness in aging and disease.

## Supporting information

Supplementary_movie_1_mNGCol_particles

Supplementary_movie_2_HT1080_mNGcol_Trackmate

Supplementary_movie_3_HT1080_mNGcol_withMTs

Supplementary_movie_4_secretion

Supplementary_movie_5_Soas2_mNGcol_Trackmate

Supplementary_movie_6_SOAS2_mNGcol_TrackMate_merged_with_MT_labelling

Supplementary_movie_7_SOAS2_mNGcol_TrackMate_merged_with_MT_labelling

Supplementary_movie_8_fibril_bleaching

Supplementary_movie_9_fibril_growth

Supplementary_movie_10_collagen_growthalongfibril_bundling

Supplementary_movie_11_bifurcation_bundling_zippering

Supplementary_movie_12_Collagen_crossovers_with_maxima

Supplementary_movie_13_fibril_meeting

Supplementary_movie_14_tensioning

Supplementary_movie_15_loopbackandincrease

Supplementary_movie_16_looping

Supplementary_movie_17_Interwinding

## Acknowledgements

This research was funded by Procter & Gamble. We would like to thank Joanne Robson (Durham University) for technical assistance with microscopy, Dr Adrian Brown (Durham University) for help with proteomic analysis, Dr Andrew Iskauskas and Prof Toby Breckon (Durham University) for helpful discussions on microscopy data analysis and Dr John Oblong, Dr Brad Jarrold and Dr Teresa DiColandrea (Procter and Gamble) for helpful discussions about the project.

## Author contributions

T.J.H, A.M.B, A.M, M.F, M.E conceived the project and T.J.H, A.M.B, A.M supervised research, T.J.H, A.M.B, A.M., O.K, E.C, M.B designed methodology. O.K, E.C, S.B. M.B., performed research and with T.J.H analysed data. T.J.H, A.M.B, A.M, E.C and O.K wrote the paper with edits from M.F., M.E, M.B.

## Methods

### Cell lines

The human fibrosarcoma cell line HT1080 (ATCC, CCL-121^TM^) was maintained in Dulbecco’s modified Eagle’s medium (DMEM). The primary human osteogenic sarcoma cell line Saos-2 (ECACC, 89050205) was maintained in McCoyʹs 5A modified medium. Dermal BJ Fibroblasts (gifted by Procter& Gamble) were maintained in minimum Eagle’s medium (MEM). All cell lines were supplemented with 10% foetal bovine serum (FBS), 100 µgml^-1^ penicillin, 100 µgml^-1^ streptomycin and 2 mM glutaMAX and were maintained at 37 °C and 5% CO_2_. Cells were sub-cultured twice weekly.

### Transfection of cells with mNG-COL1α2

The GFP-Col1α2 published by Dallas et al. (2018) was modified to substitute mNG as the fluorescent protein tag and include the CMV promoter to drive transcription. The construct was synthesized by VectorBuilder (VectoBuilder.com) using the pRP[Exp] vector backbone (Kaufman, R.J. 2000^40^). HT1080 cells were transfected using the jetPEI DNA transfection reagent (Polyplus-transfection SA) when cells reached 50-70% confluency. A 4:1 ratio of jetPEI to plasmid DNA (w/w) was diluted in NaCl following the manufacturer’s instructions and added to DMEM growth media. Imaging was conducted 24 hours post-transfection._Saos-2 and BJ fibroblast cells were transfected using Lipofectamine 3000 Reagent (Invitrogen, ThermoFisherScientific) when cells reached 70-90% confluency. For Saos-2, a 3:3:2 ratio (v/w/w) of Lipofectamine 3000 reagent, P3000 reagent, and plasmid DNA were diluted in OptiMEM following the manufacturers’ instructions and added to McCoyʹs 5A Medium. If collagen deposition was required, cells were supplemented with 20 μM ascorbic acid immediately after transfection. For BJ fibroblasts the same component ratio and dilution was used, however the transfection was conducted in the cell suspension phase, with cells at 80% confluency. For collagen deposition cells were supplemented with 200 μM ascorbic acid immediately after transfection. For all cells Imaging was conducted 48 hours post-transfection.

### Imaging

Live-cell confocal images were captured using a Zeiss 880 with Airyscan, oil 63x Plan Apochromat DIC II 1.4NA lens, Zeiss Zen 2.3 SP1 FP3 (Black) V. 14.0.22.201. Laser excitation 488nm emission BP 495-550 for mNGCol1alpha2, Laser excitation 594 emission LP 570 for AntiCol1/Alexa 594 and Laser excitation 633 emission-LP 645 for SiR-tubulin. Long term timelapse movies were captured using an Andor Revolution XD Spinning Disk with a 100x oil UPlanSApo1.4NA lens, Andor IQ V3. Laser excitation 488nm, emission 525/30nm. 3D-SIM images were captured using a GE Healthcare/Deltavision OMX V4 with an Olympus PlanApoN 60x oil 1.42NA. TIRF images were captured using a Deltavision OMX V4 with an Olympus TIRF 60x ApoN 1.49NA / Olympus TIRF 100x UApoN 1.49NA. Both 3D-SIM and TIRF Laser excitation 488nm emission 528/48nm. For all live-cell imaging experiments environmental controls were set to 37°C and 5% CO_2_

### Stains and treatments

Microtubules were visualised by staining with 1μM SiR-tubulin (Tebubio, SC002) for 30 min prior to imaging. BFA treatment was conducted at 100 ngml^-1^ for 60 min prior to fixation.

### Image analysis

#### Vesicle analysis

Analysis was conducted using Fiji. Images were thresholded to remove background signal and the TrackMate plugin (Tinevez et al., 2017^41^) was used to ascertain total number of collagen vesicles, and mean velocity and displacement data.

#### Co-localization

The Fiji plugin Coloc 2 was used to perform Coste’s regression to generate a 2D intensity histogram, calculate a Person’s coefficient and perform Coste’s significance test between the red and green channels of interest (Costes et al., 2004^42^).

### Immunofluorescence sample preparation and staining

Cells were seeded on 18 mm No. 1.5 coverslips and transiently transfected with mNGCol1α2 by the method listed above. Following any treatments, cells were fixed in 4% PFA for 10 min and the membrane was permeabilised with 0.1% Triton X-100 for 10 min. Blocking was conducted in 2% filtered BSA diluted in PBS for 30 minutes. Cells were incubated with primary antibody (1:200 in 2% BSA in PBS) overnight, and secondary antibody for 1 hr (1:500 in 2% BSA in PBS) Cells were stained with 40 ng/ml DAPI for ten minutes before mounting in Vectashield. Col1 was stained with goat anti-type1 collagen-UNLB (Southern Biotech, 1310-01), and P4HB was stained with mouse anti-P4HB [RL90](AB2792 Abcam) The secondary antibodies were donkey anti-goat conjugated to Alexa Fluor Plus 594 (ThermoFisher Scientific, A32758) or donkey anti-mouse conjugated to Alexa Fluor 594 (Thermofisher Scientific, A21203) respectively.

### Immunoprecipitation and mass spectroscopy

HT1080 cells were lysed in radioimmunoprecipitation assay (RIPA) buffer for 5 min (1% v/v Triton X-100), 50 mM Tris HCl, pH 8, 150 mM NaCl, 0.5% w/v Na-deoxycholate, 0.1% w/v sodium dodecyl sulphate (SDS), 1X PI protease inhibitors and 1x phosSTOP (Roche). Lysates were centrifuged at 16,100g at 4 °C for 10 min. Post-nuclear supernatants were subject to IP. A 30% suspension of protein A sepharose beads in RIPA buffer were incubated with the mNG antibody (32F6 Chromotek) for 1 hr at 4 °C. The supernatant was removed, and beads washed 3x with RIPA buffer for 5 min. Cell lysate was mixed with the beads for 1 hr at 4 °C. After incubation the beads were centrifuged at 6,000 g, the supernatant discarded, and the remaining beads washed 5 times in RIPA buffer. The beads were resuspended in elution buffer (50 mM NH4HCO3, 50 mM DTT, 1% SDS) to elute the mNGCol1α2 and its interacting proteins. Proteins were digested with filter aided sample preparation (FASP) and analysed by mass spectrometry. Sample fractions (5 μg of peptides) were analysed using an ekspertTM nanoLC 425 with low micro gradient flow module (Eksigent) attached to a quadrupole Time-Of-Flight (QTOF) mass analyser (TripleTOF 6600, SCIEX) connected to a DuoSpray source (SCIEX) and a 50-micron ESI electrode (Eksigent). The samples were loaded and then washed on a TriArt C18 Capillary guard column 1/32”, 5μm, 5 x 0.5mm trap column (YMC). Chromatographic separation was performed over 57 min on a TriArt C18 Capillary column 1/32”, 12 nm, S-3 μm, 150 x 0.3 mm (YMC) at a flow rate of 5 μl min-1 with a linear gradient of 3-32% acetonitrile, 0.1% formic acid over 43 min; then to 80% acetonitrile, 0.1% formic acid over 2 min, held for 3 min before returning to 3% acetonitrile, 0.1% formic acid and re-equilibrated. Analyst software (version 1.7.1, Applied Biosystems) was used to acquire the MS and MS/MS data.

