## Supplemental Information for "New imaging tools reveal live cellular collagen secretion, fibril dynamics and network organisation"

**Kent *et al.* 2024**

**Supplementary Figure 1:** Assessment of the binding partners, processing and trafficking of the mNG-Col1α2 fusion protein.

**A**

| Gene Name | Peptides | % Coverage | GO function |
|---|---|---|---|
| mNGCol1α2 | 28 | 26 | Collagen fusion Protein |
| PDIA1 | 6 | 15 | Protein folding |
| P4HB | 4 | 10 | Collagen fibril organisation |

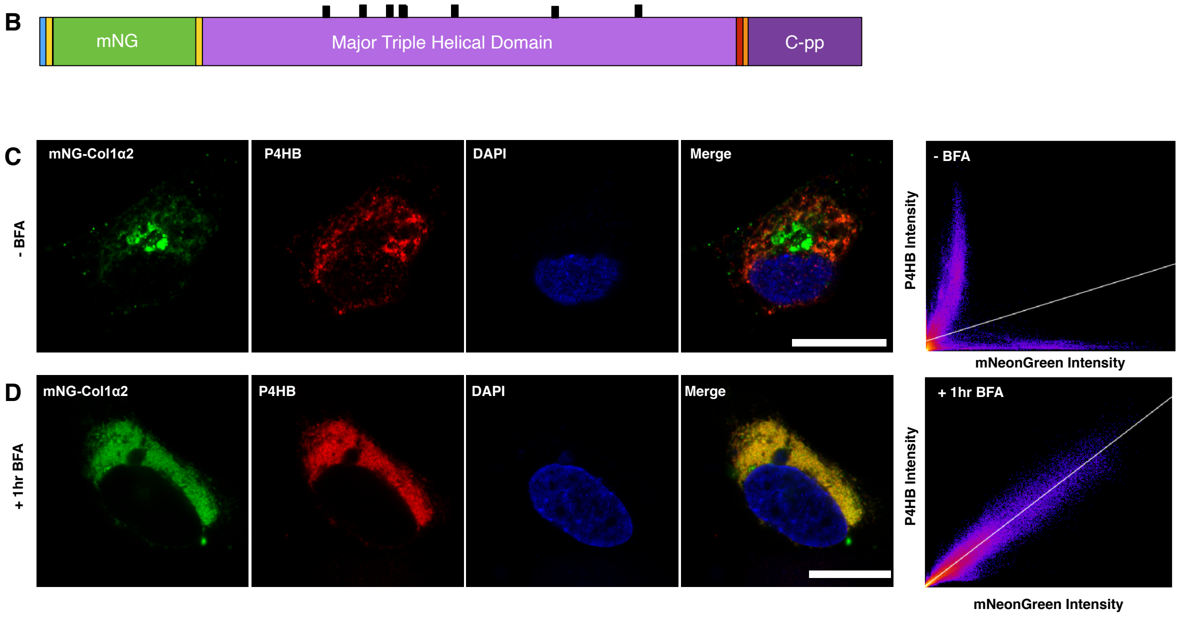


**(A)** Table of mNG-Col1α 2 interacting proteins identified by mNG IP followed by DDS MS. **(B)** Schematic diagram showing the location of hydroxylation within the mNG-Col1α 2 polypeptide as analysed by mNG IP followed by DDS MS. **(C)**Immunofluorescence confocal imaging of mNG-Col1α2 (green) expressing HT1080 cell, co-labeled for the ER marker and collagen processing chaperone P4HB (red) and DAPI. **(D)** Same staining conditions as C but following BFA treatment. Following BFA treatment the mNG-Col1α2 signal is retained in the ER. To the right are graphical representations of the Pearsons colocalization correlation for each image set. Scale = 20 μm.

**Supplementary Figure 2:** BFA treatment of Saos-2 cells expressing the mNG-Col1α 2 fusion protein

**(A)** Immunofluorescence confocal imaging of mNG-Col1α 2 (green) expressing Saos-2 cell, co-labeled for the ER marker and collagen processing chaperone P4HB (red) and DAPI. **(B)** Same staining conditions as A but following BFA treatment. Following BFA treatment the mNG-Col1α2 signal is retained in the ER. Scale=10μm

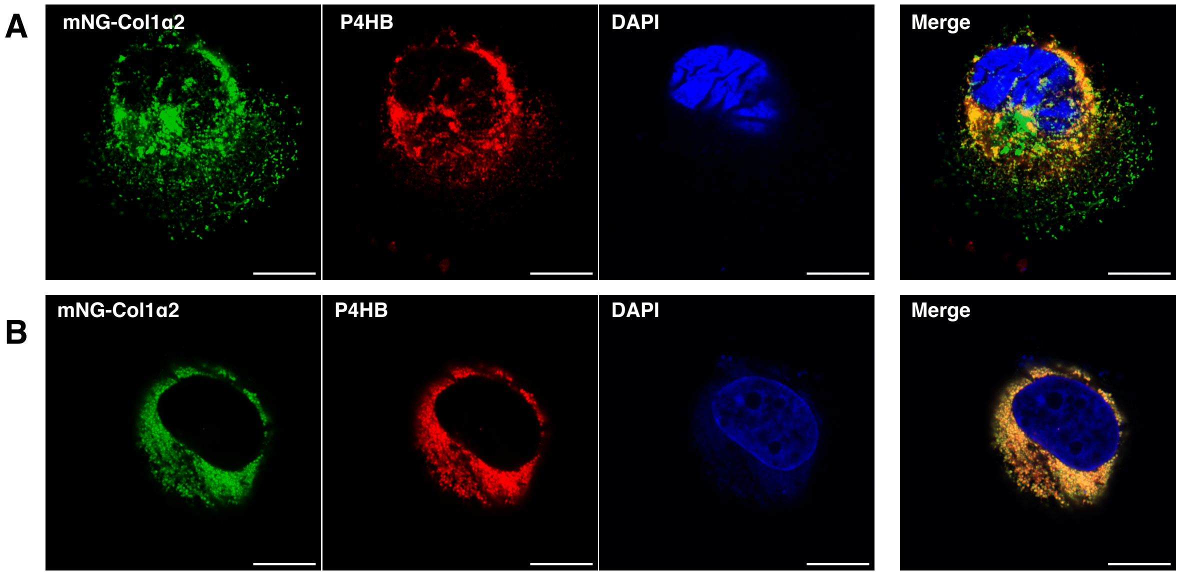


**Supplementary Figure 3: BJ Fibroblast cells expressing the mNG-Col1α2 fusion protein.** Two example cells show motile collagen carriers and deposit mNG-Col1α2 containing collagen fibrils, comparable to those seen in Saos2 cells. Scale = 20 μm

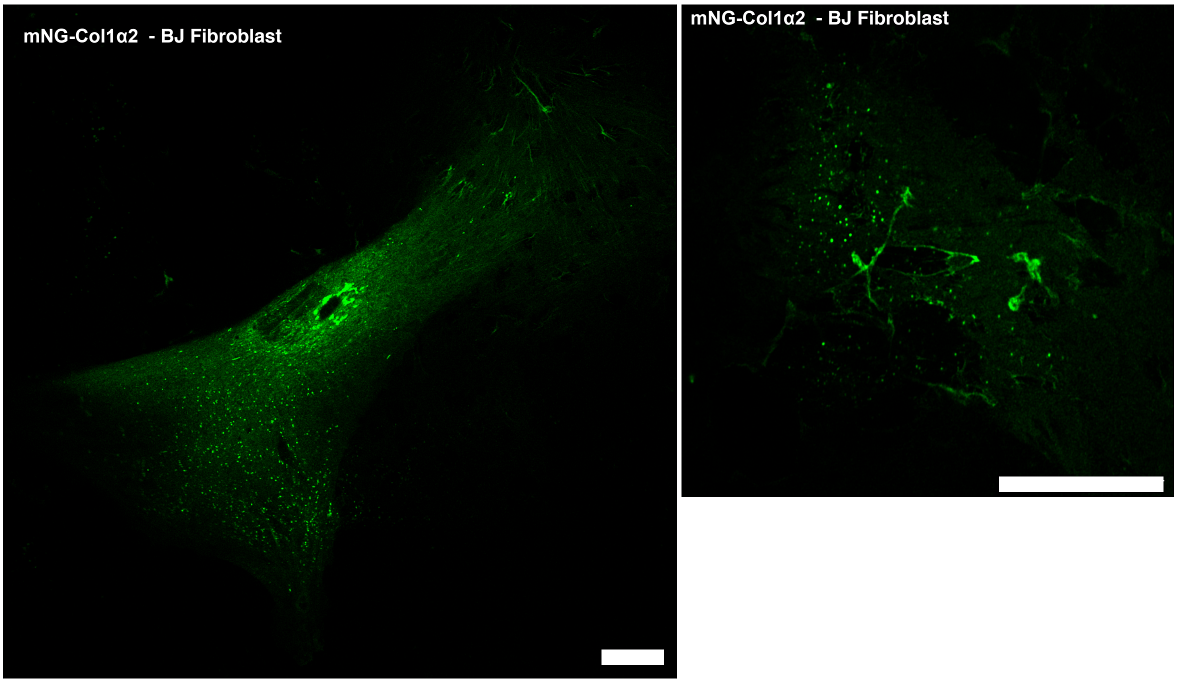


**Supplementary Movie 1: Movement of mNG-Col1alpha2 containing carriers in HT1080 cells.** Analysis of mNG-Col1α2 labelled collagen carriers which move at high speeds, predominately radiating towards the cell periphery. The movie was taken at 4.84 fps, with a total of 160 frames.

**Supplementary Movie 2: Trackmate analysis of mNG-Col1αa2 carriers.** Video of mNG-Col1α2 containing carriers in HT1080 cells (as in supplementary movie 1) with Trackmate tracks overlaid. Trackmate analysis enabled visualisation of carrier paths and quantification of individual velocities. The movie was taken at 4.84 fps, with a total of 160 frames.

**Supplementary Movie 3:** mNG-Col1α2 containing carriers associate with and travel along microtubules. The movie was taken at 1 frame every 8.66 seconds for a total of 10 frames.

**Supplementary Movie 4: mNG-Col1$\alpha$2 secretion from HT1080 cells.** Video of a single mNG-Col1$\alpha$2 fusion protein containing carrier. Movement occurs towards the PM where it remains stationary before fusing and releasing contents. Frame rate 10 ms per frame or 100 fps.

**Supplementary Movie 5:** Video of the movement of mNG-Col1$\alpha$2 containing carriers in Saos-2 cells with Trackmate tracks overlaid. The movie was taken at 4.84 fps, with a total of 160 frames.

**Supplementary Movie 6: mNG-Col1$\alpha$2 particles in Saos-2 cells move along microtubules.** Video of mNG-Col1$\alpha$2 containing carriers in Saos2 cells co-stained for microtubules with SiR tubulin. Trackmate tracks of each carrier are overlaid. The movie was taken at 4.84 fps, with a total of 160 frames.

**Supplementary Movie 7: mNG-Col1$\alpha$2 particles in Saos-2 cells move along microtubules.** A second example Video of mNG-Col1$\alpha$2 containing carriers in Saos-2 cells co-stained for microtubules with SiR tubulin. Trackmate tracks of each carrier are overlaid. The movie was taken at 4.84 fps, with a total of 160 frames.

**Supplementary Movie 8: mNG-Col1$\alpha$2 fibril bleaching.** When a mNG-Col1$\alpha$2 fibril is exposed to continuous laser exposure the signal does not bleach uniformly, reflecting the number of mNG-Col1$\alpha$2 proteins and their distribution within the fibril. 10 frames per minute.

**Supplementary Movie 9: Collagen fibril linear growth.** Video of the linear and directional growth of a mNG-Col1$\alpha$2 containing collagen fibril. Frame rate is 1 frame every 5 minutes or 12 frames per hour.

**Supplementary Movie 10: Collagen fibril growth along an existing fibril.** This video shows an initial faint collagen fibril grow and enter the frame. There are two clear growths of a new fibril along the pre-existing fibril or path. As each fibril extends the mNG-Col1$\alpha$2 intensity increases in a directional manner. Frame rate is 1 frame every 5 minutes or 12 frames per hour.

**Supplementary Movie 11: Collagen fibril bifurcation, zippering and bundling.** Video shows a collagen fibril grow from top to bottom of the frame. In later time points this singular fibril is seen to split into two thinner fibrils which then continue to grow in the original direction. In the centre of the frame two fibrils grow in opposite directions before meeting and then bundling together. They zipper up from the bottom of the frame upwards. Finally, the remaining fibril to the right snaps across in just 4 frames to join with the fibril on the left. Frame rate is 1 frame every 5 minutes or 12 frames per hour.

**Supplementary Movie 12: Collagen fibril cross overs and fluorescence intensity maxima.** In this video several collagen fibrils grow across the frame in different directions. Dark punctate maxima of mNG-Col1$\alpha$2 signal can be seen developing on a fibril at the approximate location at which another fibril will cross over it. Once these crossover points or nodes are established the mNG-Col1$\alpha$2 maxima continue to grow in size and intensity. Frame rate is 1 frame every 5 minutes or 12 frames per hour.

**Supplementary Movie 13: Collagen fibril meeting.** In the video two. mNG-Col1α2 containing collagen fibrils grow in opposite directions towards each other. As they meet they continue to grow in their original direction but now along the opposing fibril. Frame rate is 1 frame every 5 minutes or 12 frames per hour.

**Supplementary Movie 14: Collagen fibril tensioning**. In this video a large curve of collagen straightens into a square U shape. The fibril is attached at two points and is stretched between them straightening the curve to a series of straight lines with even right angled corners. The movie is an example of a live cell imparting mechanical force on its surrounding collagen network. Frame rate is 1 frame every 5 minutes or 12 frames per hour.

**Supplementary Movie 15: Collagen fibril growth into loops.** In this movie an mNG-Col1α2 containing fibril grows and curls back on itself generating a loop. This loop then increases in thinness and intensity as new collagen fibrils grow along this looped fibril/path. Frame rate is 1 frame every 5 minutes or 12 frames per hour.

**Supplementary Movie 16: Collagen fibril looping**. In this video an existing collagen fibril is moved by the cell or flow of culture media to loop up around itself. Frame rate is 1 frame every 5 minutes or 12 frames per hour.

**Supplementary Movie 17: Collagen fibril interwinding.** This video shows two separate collagen fibrils growing along paths that wrap around each other and interwind the two strands. Frame rate is 1 frame every 5 minutes or 12 frames per hour.
